# Experimentally induced pain does not influence updating of peripersonal space and body representations following tool-use

**DOI:** 10.1101/500587

**Authors:** Axel D Vittersø, Monika Halicka, Gavin Buckingham, Michael J Proulx, Mark Wilson, Janet H Bultitude

## Abstract

Representations of the body and peripersonal space can be distorted for people with some chronic pain conditions. Experimental pain induction can give rise to similar, but transient distortions in healthy individuals. However, spatial and bodily representations are dynamic, and constantly update as we interact with objects in our environment. It is unclear whether induced pain disrupts the mechanisms involved in updating these representations. In the present study, we sought to investigate the effect of induced pain on the updating of peripersonal space and body representations during and following tool-use. We compared performance under three conditions (pain, active placebo, neutral) on a visuotactile crossmodal congruency task and a tactile distance judgement task to measure updating of peripersonal space and body representations, respectively. We induced pain by applying 1% capsaicin cream to the arm, and for placebo we used a gel that induced non-painful warming. Consistent with previous findings, the difference in crossmodal interference from visual distractors in the same compared to opposite visual field to the tactile target was less when tools were crossed than uncrossed. This suggests an extension of peripersonal space to incorporate the tips of the tools. Also consistent with previous findings, estimates of the felt distance between two points (tactile distance judgements) decreased after active tool-use. In contrast to our predictions, however, we found no evidence that pain interfered with performance on either task when compared to the control conditions. This suggests that the updating of peripersonal space and body representations is not disrupted by induced pain. Therefore, acute pain does not account for the distorted representations of the body and peripersonal space that can endure in people with chronic pain conditions.

## Introduction

The multisensory representations of our body and its surrounding space are constantly updated as we interact with objects in our environment. Work with macaques identified bimodal neurons that responded to both somatosensory and visual information near and on the hand, whose receptive fields were malleable as a function of active tool-use [1]. When monkeys actively used a rake to retrieve food, the receptive fields of these neurons expanded to include the area near to and occupied by the rake. Subsequent research in humans has shown that responses to visual, tactile, and auditory stimuli that originate near and on tools are modulated by active tool-use. Changes that arise from active tool-use are thought to reflect that the cortical representations of the body and its surrounding space have been updated to accommodate the new properties offered by the tool (for reviews see [2–4]).

Active tool-use, and the changes it causes in multisensory processing, have been used to study the representations of both the body and peripersonal space. Here we define body representation as the mental model of the body, based on proprioceptive and sensory information about the body’s state [5], for reviews, see [6, 7]. This representation is flexible, and a small degree of distortion has been demonstrated in normal cognition [8]. Peripersonal space is defined as the areas that directly surround the body that we can act upon [9], and that can contain objects that we may need to react to. Peripersonal space has been characterised by neurophysiological and neuropsychological studies with animals and clinical and non-clinical human populations (for reviews, see [10, 11]). The body and its representation are the centre of peripersonal space [12], hence these representations are to some degree related, and active tool-use can influence both. For instance, Canzoneri and colleagues [13] used an audio-tactile interaction task to assess peripersonal space, and a landmark task and tactile distance perception to examine body representation, before and after participants used a tool to retrieve distant objects. The results showed that following tool use participants perceived their arm as narrower and longer, and representations of peripersonal space surrounding the arms were extended along the axis of the tool. Therefore, active tool-use provides an opportunity to study how body and peripersonal space representations are updated.

Representations of the body and space can be distorted in people with neurological disorders like asomatognosia [14] and hemispatial neglect [15–17]. Body representation is also distorted in people with certain types of chronic pain (for reviews see [18–20]). For instance, people with chronic back pain often report a distorted sense of the size of their body near their painful area, or that parts of the body feel like they are missing [21]. Similarly, people with Complex Regional Pain Syndrome (CRPS) can report difficulties locating and recognising their affected limb, show a distorted perception of its size, and have difficulties locating touch on their affected hand (e.g. [22–29]). Distorted spatial representations have also been reported in people with pathological pain conditions. For instance, people with unilateral hand amputations underestimate the size of near space on the side of their amputation, compared to the contralateral side [30]. In CRPS patients, biases in visual and tactile attention away from the affected hand have been identified by asking patients to judge the temporal order of pairs of tactile stimuli delivered to, or visual stimuli projected near to or onto, the hands [31–34]. Estimates of the point in space that is straight ahead of the body midline made in complete darkness, thought to reflect the division between left and right space in an egocentric reference frame [35], are also deviated in people with CRPS. Such deviations in spatial perception have been reported in the direction of the affected side [36–38], and leftwards irrespectively of the affected side [39], although not all studies find evidence of deviations [40–42]. Furthermore, Sumitani and colleagues [36] found that the deviation towards the CRPS-affected side was reduced following pain reduction through a using a nerve block. Taken together, these studies demonstrate that bodily and spatial representations can be distorted in pathological pain conditions. What is unclear, however, is whether pain (or associated factors such as immobility and disuse) *precede*, or *follow* (e.g. [43]), these altered representations.

Research has started to investigate the effect of pain on spatial perception and the representation of the body in normal cognition. Pain itself is a sensory and affective phenomenon [44] that can be shaped by multisensory experiences [45] and convey spatial information about the body [19]. After participants were subjected to painful heat stimulation on one hand, their subjective body midline shifted towards the painful side, whereas vibrotactile stimulation had the opposite effect [46]. This suggests that pain can modify spatial perception in ways that cannot sufficiently be explained by tactile stimulation or attentional cueing effects. To date, only one study looked directly at how pain might alter the representation of the body in healthy subjects. Gandevia and Phegan [47] found that participants reported an average of 10% increase in the perceived size of their thumb after it had been subject to painful cooling. These studies suggest that pain might alter the representations of the body and its surrounding space, however the evidence is limited. Furthermore, to the best of our knowledge, no study has investigated the effect of pain on the ways that peripersonal space and body representation update due to experiences such as tool use.

Our study aimed to investigate the effect of induced pain on the updating of peripersonal space and body representations of healthy individuals during and following tool-use. Over three separate sessions, participants completed a tool-use task while experiencing capsaicin-induced pain in their dominant arm, and in two control conditions: active placebo and neutral (i.e., no sensory manipulation). We hypothesised that inducing pain in an arm would impair participants’ ability to update peripersonal space and body representations relative to the other conditions. We used a crossmodal congruency task (CCT) and tactile distance judgements (TDJs) to measure updating of peripersonal space and body representation, respectively.

The CCT has been used previously to investigate the effects of active tool-use on spatial representations [3, 4]. In this task, participants make judgements about vibrotactile targets presented to the hands through the handles of the crossed or uncrossed tools while ignoring visual or auditory distractors presented at the tips of the tools. After the participants have used the tools for a period of time, distractors on the same side of space as the targets typically have a larger effect on increasing reaction times and/or error rates when the tools are uncrossed than when the tools are crossed [48]. In contrast, distractors on the opposite side of space as the targets have a larger effect on performance when the tools are crossed than when the tools are uncrossed. That is, after tool use, distractors have a greater interference effect when they originate from the same *tool* as targets, rather than from the same *side of space* as the target. Maravita and colleagues [48] interpreted this pattern to indicate a change in peripersonal space representations, which was further suggested by the fact that they showed the pattern of interference developed gradually over a period of ongoing tool use (although see Holmes [2012] for an alternative interpretation), but did not develop when tools were held passively for the same period of time and number of trials. We predicted that pain would interfere with the emergence of tool-specific effects of distractors on judgements made about the targets. That is, we expected to see a weaker interaction between the arrangement of the tools, the visual field in which visual distractors appear relative to vibrotactile targets, and the vertical congruence of visual distractors relative to vibrotactile targets for the pain condition, relative to the two control conditions.

TDJs have been used to measure updating of body representation following tool use. Distances between two touched locations on the arm that are oriented parallel to the axis of the tool are perceived to be shorter after active tool-use. This is thought to indicate that body representation is altered by tool use, such that the forearm is perceived to be longer [13, 50–53]. We predicted that pain would interfere with the way in which tool use altered these TDJs, such that distance estimates would have a smaller decrease when participants were in pain.

## Methods

### Design

We used a repeated-measures design with three sessions, corresponding to three sensory conditions: Pain, active placebo, and neutral (i.e., no sensory manipulation). We used a Crossmodal Congruency Task (CCT) adapted from Maravita and colleagues [48] to measure changes in peripersonal space and Tactile Distance judgements (TDJs) adapted from Canzoneri and colleagues [13], Miller and colleagues [50, 52], Longo and Haggard [51], and Taylor-Clarke and colleagues [53] to measure changes in body representation. We wanted to know if the effects of unilateral pain induction would be specific to the stimulated arm, or global (i.e. extend to the unstimulated arm), by comparing CCT and TDJ performance between the two arms. In addition to these tasks, we also used several measures to monitor the sensory and cognitive effects of the sensory manipulations. We asked participants to give numerical ratings of pain intensity. We also used sensory testing (Mechanical Pain Threshold [MPT], Mechanical Detection Threshold [MDT], Two Point Discrimination Threshold [TPD]), and questionnaire measures (The Bath CRPS Body Perception Disturbance Scale [BPD; [54]], the Short-Form McGill Pain Questionnaire 2 [SF-MPQ-2; [55]]), to characterise any secondary changes caused by our sensory manipulation. The protocol was preregistered on the Open Science Framework (https://osf.io/8fduw/register/565fb3678c5e4a66b5582f67).

### Participants

Thirty-one participants completed the study tasks under three sensory conditions (pain, active placebo, and neutral) in a randomized, counterbalanced order. One person was excluded because she did not report any pain (0/10 on a Numerical Rating Scale [NRS]) for 40 minutes following the application of capsaicin, even when we attempted this condition on a second occasion. One person repeated the pain condition (their session 1) due to low pain ratings in the first attempt that was completed (*M* NRS after the sensory manipulation period during the first attempt 0.6/10, *SD* = 0.97 and the second attempt 1.4/10, *SD* 0.70). One person repeated the neutral condition (their session due to equipment malfunctioning during the CCT. The repeated sessions were completed in full, and took place on a different day. The mean age of the final sample was 21.6 years (*SD* = 4.3), of which 22 (73.3%) were women. Two participants were left-handed (*M* = -70.0, *SD* = 14.1), one ambidextrous (score of 30), and the remaining 27 were right-handed (*M* = 83.6, *SD* = 17.7), as indicated by the Edinburgh Handedness Inventory [56], in which extreme left and right handedness is indicated by scores of -100 and 100, and scores between 40 and -40 indicate ambidextrousness. All participants reported having normal or corrected to normal vision, and that they did not have a chronic pain condition. Participants with self-reported sensitive skin, epilepsy, high blood pressure, recent heart problems, a history of stroke, vascular problems, an allergy to capsaicin, or who were pregnant or breastfeeding were excluded to satisfy local safety guidelines for the use of capsaicin cream. Participants signed consent and safety forms prior to participating, and consented for their data to be used upon completion of the study. The study adhered to the 2013 Declaration of Helsinki, and received approval from the local ethics committee. Participants received £30 for their involvement.

### Materials

For the pain condition, a 1cm wide band of Ungentum cream infused with a 1% concentration of capsaicin (the Specials Laboratory, United Kingdom), amounting to approximately 5g of cream, was applied to the dominant arm, just proximal to the elbow. For the ambidextrous person, the cream was applied to the right arm because this was the participant’s self-reported dominant side. The cream was contained within two bands of microporous tape, and covered with cling film. This was fitted so that participants could flex and extend their elbow with ease, so as not to impede their ability to manoeuvre the tools. Applying a band of capsaicin that reaches around the arm in this location generates a burning pain that penetrates into the arm [57], and is accompanied by cutaneous vasodilation and hyperalgesia [58]. This method has been used previously [57, 59]. For the active placebo, a ‘warm-up’ gel (Elite Ozone^®^) was used to create a non-painful warming sensation. The site and application procedures were identical to that of the capsaicin. No cream was applied in the neutral condition.

The final design of the CCT was informed by pilot research (n = 42). The materials used were based on the study by Maravita and colleagues [48]. Two 75cm long tools (see Fig 1) that resembled golf clubs were constructed from aluminium. Two red Light Emitting Diodes (LEDs) were embedded in the distal end of each tool. Two electromagnetic solenoid-type stimulators (Tactor Minature Stimulators, Dancer Design, United Kingdom) were embedded in each of the handles to deliver vibrotactile stimulation. The LEDs and vibrotactile stimulators were controlled by a 4-channel amplifier (TactAmp 4.2, Dancer Design, United Kingdom) operated by Matlab 2014b (MathWorks). On each tool, one LED and one vibrotactile stimulator was positioned above the central axis of the tool, and one LED and one vibrotactile stimulator below it. The tools had wooden pegs attached vertically to their far ends, near the LEDs, in the ‘blades’ of the tools. The pegs slotted into holes in a wooden board (80 x 100 cm) that were 15 cm away from the distal end of the board, and 15 cm to the left and right from the central axis of the board. This ensured that the ends of the tools were always placed in the same position regardless of whether the tools were crossed or uncrossed. A fixation light was located at the central axis of the board, 15cm from the distal end. A 5 cm wide blue mark was placed on the handle of each tool 30 cm away from the distal end to indicate points at which participants should cross the tools. Two triple switch foot pedals (Scythe, USA) with custom software were used to collect participants’ responses. White noise was played on headphones to mask any sound of the vibrotactile stimulation. A chinrest was used to ensure that participant’s heads remained in a consistent position. Two webcams were positioned in line with participant’s sagittal plane, at the end of and 20 cm away from the board, so that the experimenter could monitor gaze throughout the task, and record participant’s movements for offline evaluation of movement quality.

**Fig 1.**
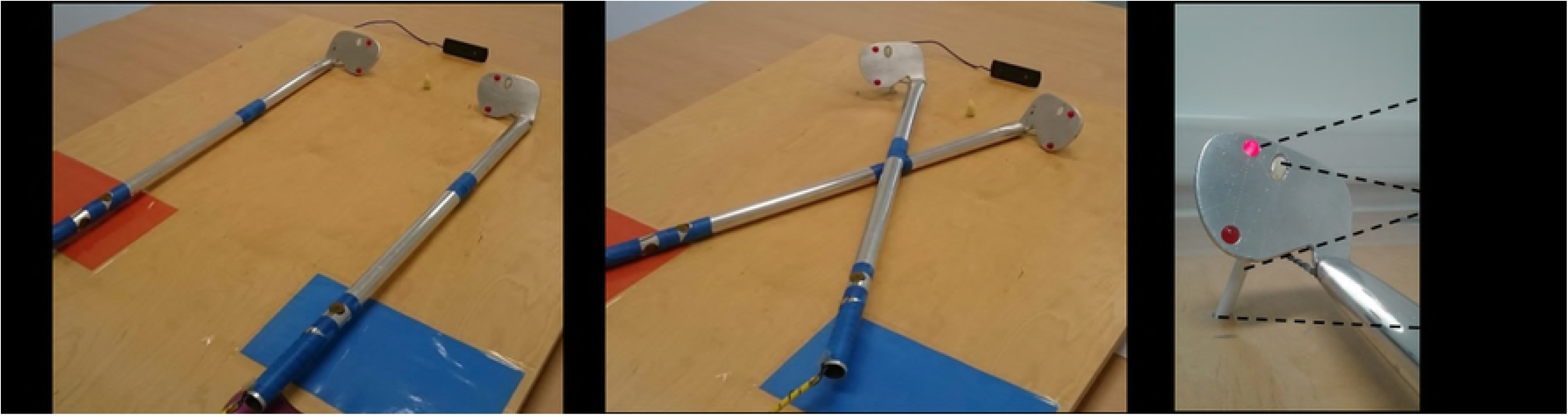
Tool-use materials. From left to right, the images depict the uncrossed (A), and crossed (B) tools, and a close-up of the end of a tool (C). The tools have red Light Emitting Diodes (LEDs) embedded at the far ends of the tools, and vibrotactile stimulators embedded in the handles. A vertical peg was attached to the far ends of the tools (white oval), which slotted into holes in the wooden board to ensure the position of the tips of the tools was consistent for crossed and uncrossed trials (C). An off-white LED fixation point was positioned with equal distance to the ends of both tools. A webcam was placed in line with the fixation light and the chinrest, which were aligned with participants’ sagittal plane.

Two flat-ended circular rods (1 mm diameter) were used for tactile distance judgements (TDJs). They were attached to a bow compass, so that the distance between them could be accurately adjusted. MPT was assessed using seven pinprick stimulators (MRC Systems GmbH, Germany), ranging from 8 mN to 512 mN in force. Twenty Von Frey Filaments were used to measure MDT (BioSeb, France), ranging from 0.008 g to 300 g in weight. TPD was quantified using an Exacta™ two-point discriminator (North Coast Medical, USA), ranging from distances of 2 mm to 20 mm in distance. A handheld infrared thermometer with an 8:1 distance to spot size ratio, and a red laser aim was used to measure the temperature of participants’ hands.

The BPD [54] is an unvalidated 7-item questionnaire designed to characterise distorted body perception in CRPS. In includes questions about the awareness of, attention to, emotional valance of, and desire to amputate the affected area. For this study, participants were instructed to answer about the stimulated (dominant) arm. Scores can range from 0 to 57, where a higher score indicates greater distorted body perception.

The SF-MPQ-2 [55] is a 24-item questionnaire assessing the symptoms of neuropathic and non-neuropathic pain. Participants rate the intensity of their pain for each of 22 pain qualities (e.g. sharp, aching, hot-burning) on a scale of 0 (‘none’) to 10 (‘worst possible’). The SF-MPQ-2 has been validated for acute pain populations [60].

### Procedure

Fig 2 shows an outline of the procedure for each session. Informed written consent was obtained and self-reported handedness was recorded upon commencing the first session. Then, the first set of sensory tests (i.e. MPT, MDT, TPD) were performed on the middle finger (digit 3) of the dominant and non-dominant hands. MPT, and MDT were assessed following a standardised protocol [61]. Five values for each subthreshold and suprathreshold were recorded for each sensory test. That is, we recorded when touch was detected or not for the MDT; sharp and blunt sensations for the MPT; and the distance (mm) at which two points were perceived as one or two were recorded the TPD. Then the capsaicin cream or ‘warm-up’ gel was applied for the pain and active placebo conditions, respectively. Pain ratings and dominant hand temperature were recorded every minute following cream application in the Pain and Active Placebo conditions, or upon completion of sensory testing in the Neutral condition. To allow the capsaicin to take effect, we waited until participants’ pain ratings exceeded 5/10, or until three identical consecutive ratings >2/10 were given. In the two control conditions, we waited for 15 min. Then we conducted a second set of sensory tests, and participants completed the BPD and SF-MPQ-2.

**Fig 2.**
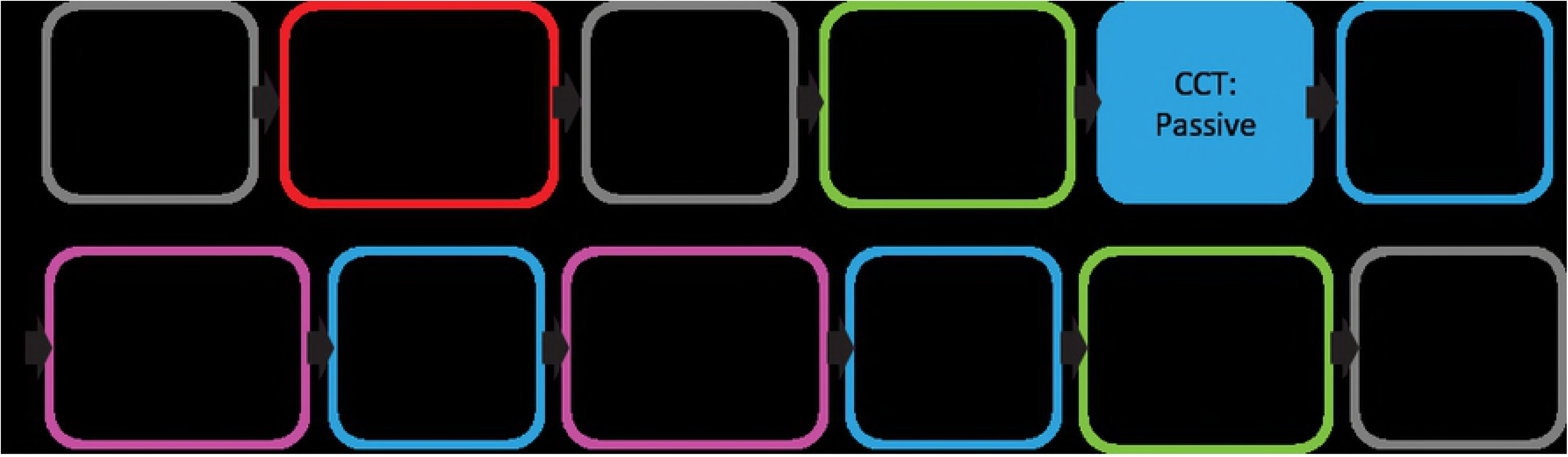
Procedure for each experimental session. During the sensory manipulation phase the experimenter applied a capsaicin cream (pain condition) or warm-up gel (active placebo condition) to the participant’s dominant arm, or there was no manipulation (neutral condition). During the passive stage (set 1) of the Crossmodal Congruency task (CCT) the experimenter changed the tools between the crossed and uncrossed positions. During the active stages of the CCT (sets 2-4) participants manoeuvred the tools themselves when changing position. The interactive tool-use task involved retrieving and sorting beanbags, using the same tools that were used for the CCT (see Fig 1).

We then administered the first TDJ task. The participant sat with their head in the chin rest with their eyes open and gripped the uncrossed tools. The experimenter applied the flat-ended circular rods to along the radial side of the participant’s forearm (i.e. proximal-distally). Three distances (4, 6, and 8 cm) were presented on one arm in a randomised counterbalanced order, with one repetition for each distance. After the three TDJ trials were completed, the procedure was repeated for the second arm. The order in which the arms were tested was randomised and counterbalanced. Participants indicated the estimated distance between the two felt points using a diagram of 22 lines of different lengths ranging from 0.6 cm to 11.5 cm, in 0.5 cm increments, presented on an A4 sheet of paper.

After the first TDJ task, participants were instructed on how to perform the CCT. On each trial, participants identified the location of three 50 ms bursts of vibrotactile stimulation delivered to the thumb or middle finger of the left or right hand from the vibrotactile stimulators embedded in the handles of the tools. Three flashes of 50 ms from the red LEDs at the ends of the tools preceded each vibrotactile stimulation by 30ms to maximise the crossmodal interference [62]. Participants judged the location of the vibrotactile stimulation as either on the thumb (upper) or finger (lower), and if it was delivered to the left or right hand. Visual distractors provided no information about the location of the target vibrotactile stimulation. Participants indicated the location of the vibrotactile stimulation by pressing foot pedals using their heel (finger; “lower”) or toes (thumb; “upper”) of their left or right foot. Participants were instructed to respond as quickly and as accurately as possible. Incorrect responses and responses slower than 3000 ms caused all four LEDs to flash three times to provide feedback to the participants.

Participants completed four Sets of the CCT. Within each set were trials in which the tools were positioned in uncrossed or crossed positions. In the first set of the CCT (the “passive” set), the experimenter changed the position of the tools half-way through while the participants kept hold of the handles, comparable to the control experiment reported by Maravita and colleagues [48]. The order of tool arrangements (crossed or uncrossed) was counterbalanced between participants. For the remaining three sets of the CCT (the “active” sets), participants actively changed the arrangement of the tools between crossed and uncrossed every four trials. The cue to change arrangement was all four LEDs illuminating. Before commencing the first (passive) CCT set, participants completed a practice set of 16 trials during which they held the tools still and uncrossed to ensure that they understood the CCT task. The practice set was repeated until the participant responded correctly on >80% of trials.

The CCT enabled the evaluation of the effect of the visual distractors on detection of vibrotactile stimulation depending on whether tools were uncrossed or crossed (Tool Arrangement), the distractor was in the same or the opposite side of space as the target (Visual Field), and the distractor was in the same or the opposite vertical elevation as the target (Congruence). All possible combinations of Tool Arrangement, Visual Field, and Congruence relative to the vibrotactile target were delivered in a random order over every 32 trials. Each set was comprised of 96 trials, giving 384 trials per session. Using this procedure, we could also examine changes in the effect of the distractors over time by comparing performance in the four sets.

Actively changing the tool arrangement was the key manipulation in Maravita and colleagues’ [48] study to elicit the changes to CCT performance generated by active tool-use. However, following piloting, we added an additional interactive tool-use tasks in-between the second and third, and third and fourth sets of the CCT to amplify the desired effect. The task consisted of approximately 5 minutes of using the tools to sort and retrieve distant beanbags, using the same equipment as for the CCT. Participants sorted beanbags by colour, and retrieved them from the distal end of the board to coloured squares (see Fig 1) on either the left of right side of the board’s proximal end. This was inspired by comparable paradigms involving active tool-use [63–65]. Upon completion of the last set of the CCT the second set of TDJs was administered, and the second set of sensory testing was conducted. Each session lasted approximately 2 hours.

In addition to the pain ratings recorded during the pain ramp-up period in the pain condition, or for 15 minutes in the active placebo and neutral conditions, participants provided an additional 12 pain ratings between different experimental tasks. They provided pain rating before each set of TDJs, and two rating for the first CCT set (passive), and then before and after each of the active CCT sets.

### Analyses

We examined error rates and reaction times (RTs) from the crossmodal congruency task in separate 3×4×2×2×2×2 repeated-measures analyses of variance (ANOVAs). The independent variables for the CCT were the Sensory Condition (pain, active placebo, neutral), Set (set 1 [passive], set 2, set 3, set 4, Side of Body on which the vibrotactile stimulation occurred (dominant, non-dominant), Tool Arrangement (uncrossed, crossed), Visual Field in which the distractor occurred relative to the target (same, opposite), and Congruence (congruent, incongruent). The median RTs and percentage of errors were calculated within level of the relevant conditions, after excluding trials with RTs <200 ms or >3000 ms. Only trials with correct responses were used to calculate the median RTs. The critical interaction that we were interested in was that between Tool Arrangement, Visual Field, and Congruence. Therefore, only interactions involving all three factors Tool Arrangement, Visual Field, and Congruence were considered relevant for addressing the aim of the study. We also considered that the three-way interaction should normally develop over time spent engaged in active tool-use (as reported by Maravita and colleagues [48]), which would result in interactions involving the four factors Set, Tool Arrangement, Visual Field, and Congruence. In their study [48], tool-use-dependent effects were only significant for RTs, not for error rates. Therefore, to be concise we will only report results from the RTs of the CCT in the main article (see S1 Text for the results of CCT error rate analyses). To aid interpretation of the results, we subtracted congruent from incongruent trials to calculate crossmodal interference for follow-up contrasts.

A mean score for the TDJs was computed for each Sensory Condition (pain, active placebo, neutral), Set (pre tool-use, post tool-use), and Side of Body (dominant, non-dominant), and analysed using a 3×2×2 repeated-measures ANOVA.

MPT, MDT, and TPD thresholds were calculated as geometric means of each value for which a participant’s responses changed (e.g. from blunt to sharp for MPT). Performance on sensory tests was analysed using ANOVAs with Sensory Condition (pain, active placebo, neutral), Set (pre manipulation, post manipulation), and Side of Body (dominant, non-dominant) as independent variables. For subsequent covariate analyses, changes on the sensory tests were calculated by subtracting pre- from post-sensory manipulation scores, within each level of Sensory Condition (pain, active placebo, neutral) for the dominant (stimulated) side of the body. The 12 pain ratings recorded after the sensory manipulation period (i.e. between sets of TDJs and CCT) were averaged across tasks, within each level of Sensory Condition, for each participant, for covariate analysis. Changes in hand temperature were calculated by subtracting the first recording (i.e. 1 minute after the sensory manipulation) from the last recording (e.g. after 15 minutes for the active placebo and neutral conditions), for each participant and within each level of Sensory Condition. To evaluate if there were any differences in participants’ movements across the three sessions that could account for any difference between the effects of tool use, movements were scored by a research assistant from video recordings taken during the first and last two minutes of the active CCT, and during the interactive tool-use task. The research assistant, who was blind to the hypotheses and task conditions, rated the speed, ease, and control of movement in each video from 0 (‘worst imaginable’) to 10 (‘best imaginable’). A mean value was calculated from the speed, ease, and control of movement for each session, within each participant. Age, the total score for the SF-MPQ-2,BPD, change in hand temperature, average pain ratings, and changes on sensory tests from pre to post sensory manipulation were explored as covariates in analyses of covariance (ANCOVAs) of the CCT and TDJ. Greenhouse-Geisser corrections were used when sphericity was not satisfied. Holm-Bonferroni corrections [66] were used for follow-up t-tests, and indicated by “*p*_adjusted_”. See preregistration (https://osf.io/8fduw/register/565fb3678c5e4a66b5582f67) for full list of planned confirmatory and exploratory analyses.

## Results

### Sensory measures

The mean duration for pain ratings to reach 5/10 or plateau after the capsaicin was administered was 16.7 minutes (*SD* = 7.62), see S2 Fig for time course. There were no differences between the changes in hand temperature as measured on the tip of digit 3 for the pain condition (*M* = -0.79°C, *SD* = 0.62), active placebo condition (*M* = -0.60°C, *SD* = 1.16), and neutral condition (*M* = -0.98°C, *SD* = 0.62) over this period *F*(1.57, 44.07) = 1.24, *p* = .292, η^2^_p_ = .04. There were no differences in the research assistant’s ratings of the movement between the pain condition (*M* = 5.73, *SD* = 1.16), active placebo condition (*M* = 5.68, *SD* = 1.20), and neutral condition (*M* = 5.60, *SD* = 1.25), *F*(2, 27) = 0.48, *p* = .627, η^2^_p_ = .03. The mean pain ratings averaged across the TDJs and CCT tasks were 5.06 (SD = 1.88) for the pain condition, 0.27 (SD = 0.46) for the active placebo condition, and 0.02 (*SD* = 0.06) for the neutral condition. Pain ratings for the pain condition were significantly higher than both the active placebo, *t*(29) = 14.16, *p*_adjusted_ < .001, *d* = 5.26, and neutral conditions, *t*(29) = 14.70, *p*_adjusted_ < .001, *d* = 5.46. Participants also reported higher pain in the active placebo condition than the neutral condition, *t*(29) = 2.84, *p*_adjusted_ = .030, *d* = 1.06. Overall, these results show the capsaicin cream induced significant pain relative to the other two conditions, without influencing movement ratings or hand temperature.

There were no changes in TPD (*M*=0.00 mm, *SD* = 0.25) or MDT (*M* = 0.00 g, *SD* = 0.01) from pre to post sensory manipulation when considered across all three sensory manipulations, or any two-way interactions of Time with Sensory Condition or Side of Body *F*s ≤ 2.99, *p*s ≥ .094, η^2^_p_ ≤ .09 (See S1 Table). For MPT there was an interaction between Sensory Condition x Time x Side of Body, F(2, 28) = 4.42, *p* = .021, η^2^_p_ = .24. This reflected that there was an increase in MPT for the dominant (stimulated) arm in the pain condition, t(29) = 2.34, *p*_adjusted_ = .048, *d* = 0.87, as MPT increased from pre (*M* = 178.3 mN, *SD* = 136.8) to post (*M* = 224.6 mN, *SD* = 161.5) the application of capsaicin cream. Follow-up analysis showed that there was a significant increase in MPT for the non-dominant side of the body in the active placebo condition from pre sensory manipulation to post CCT (*M* = 217, *SD* = 146.7), *t*(29) = 3.34, *p*_adjusted_ = .005, *d* = 1.24. There were no changes in MPT from pre to post sensory manipulation in any of the other levels of Sensory Condition by Side of Body, *t*s(29) < 1.60, *ps*_adjusted_ ≥ .214, *ds* ≤ 0.59.

Age, SF-MPQ-2 scores, BPD scores, hand temperature, average pain ratings, and changes in sensory testing were explored as covariates for the analyses of the TDJ and CCT. When analysed within each Sensory Condition, the covariates did not consistently interact with either RTs from the CCT, error rates from the CCT, or TDJ. That is, no covariate interacted significantly across each Sensory Condition for any outcome measure. Therefore, no covariates were included for further analysis.

### Crossmodal congruency task

All significant main effects and interactions for RTs are reported in S2 Table. There were main effects of Set, *F*(3, 27) = 45.10, *p* < .001, η^2^_p_ = .83, Side of Body, *F*(1, 29) = 7.97, *p* = .009, η^2^_p_ = .22, Visual Field, *F*(1, 29) = 6.69, *p* = .015, η^2^_p_ = .19, and Congruence *F*(1, 29) = 177.18, *p* < .001, η^2^_p_ = .86, on reaction times for the CCT. Reaction times became shorter for each set of the CCT (set 1 [passive]: *M* = 707.9 ms, *SD* = 101.88; set 2: *M* = 694.5 ms, *SD* = 93.40; set 3: *M* = 648.4 ms, *SD* = 92.02; set 4: *M* = 629.6 ms, *SD* = 97.49). Except for the difference between set 1 and 2 (*t*(29) = 1.57, *p*_adjusted_ < .128, *d* = 28), all follow-up comparisons showed a significant decrease in reaction time over time, *t*s(29) ≥ 3.46, *ps*_adjusted_ ≤ .004, *ds* ≥ 1.29. Participants responded faster to vibrotactile stimulation on their dominant (*M* = 660.3 ms, *SD* = 92.02) than their non-dominant (*M* = 679.9 ms, *SD* = 98.59) hand. Reaction times were shorter when visual distractors appeared in the same (*M* = 667.3 ms, *SD* = 93.66) than the opposite (*M* = 672.9 ms, *SD* = 93.66) visual field relative to vibrotactile targets. Responses were slower when visual distractors were incongruent (*M* = 693.3 ms, *SD* = 95.30) than congruent (*M* = 646.9 ms, *SD* = 92.57) with vertical vibrotactile target locations.

The most important finding with regards to our hypothesis was that no interactions of interest involving Sensory Condition, Tool Arrangement, Visual Field, and Congruence were observed, indicating that pain did not interfere with the reaction times on the CCT. Most importantly, there was no significant interaction for Sensory Condition x Tool Arrangement x Visual Field x Congruence, *F*(1.66, 48.21) = 0.80, *p* = .434, η^2^_p_ = .03.

The critical three-way interaction for testing the effects of tool use on peripersonal space, between Tool Arrangement, Visual Field, and Congruence, was significant *F*(1, 29) = 9.43, *p* = .005, η^2^_p_ = .25 (Fig 2). The follow-up analyses showed that there was a significant difference between incongruent and congruent distractors within each level of Tool Arrangement and Visual Field (see S3 Table), *t*s(29) ≥ 5.29, *ps*_adjusted_ ≤ .004, *ds* ≥ 1.96. Therefore, we calculated the crossmodal interference by subtracting congruent from incongruent scores, and compared this across each level of Tool Arrangement and Visual Field to evaluate what drove this interaction (Fig 3). For uncrossed tools, crossmodal interference was greater when the visual distractors appeared in the same (*M* = 72.2, *SD* = 29.6) than the opposite (*M* = 27.1, *SD* = 28.1) visual field to the vibrotactile targets, *t*(29) = 6.43, *p*_adjusted_ = .004, *d* = 2.39. When the tools were crossed, crossmodal interference was also greater when the visual distractors appeared in the same (*M* = 54.2, *SD* = 35.0) than the opposite (*M* = 31.3, *SD* = 21.7) visual field relative to the vibrotactile targets, *t*(29) = 3.31, *p*_adjusted_ = .009, *d* = 0.61, although the effect size was smaller than when tools were uncrossed. When visual distractors appeared in the same visual field as the vibrotactile targets, crossmodal interference was greater for the uncrossed than the crossed tools, *t*(29) = 3.42, *p*_adjusted_ = .014, *d* = 1.27. There was no difference in crossmodal interference when visual distractors appeared in the visual field opposite the vibrotactile targets, *t*(29) = 0.80, *p*_adjusted_ = .438, *d* = 0.30. These results suggest that peripersonal space representations were updated as a function of tool-use because the arrangement of the tools impacted on the RTs from the CCT. This is evidenced by the decreased effect size when comparing visual distractors appearing in the same or opposite side, giving rise to the critical three-way interaction. The results of the analysis of error rates (S1) were broadly consistent with the results of the analysis of reaction times.

**Fig 3.**
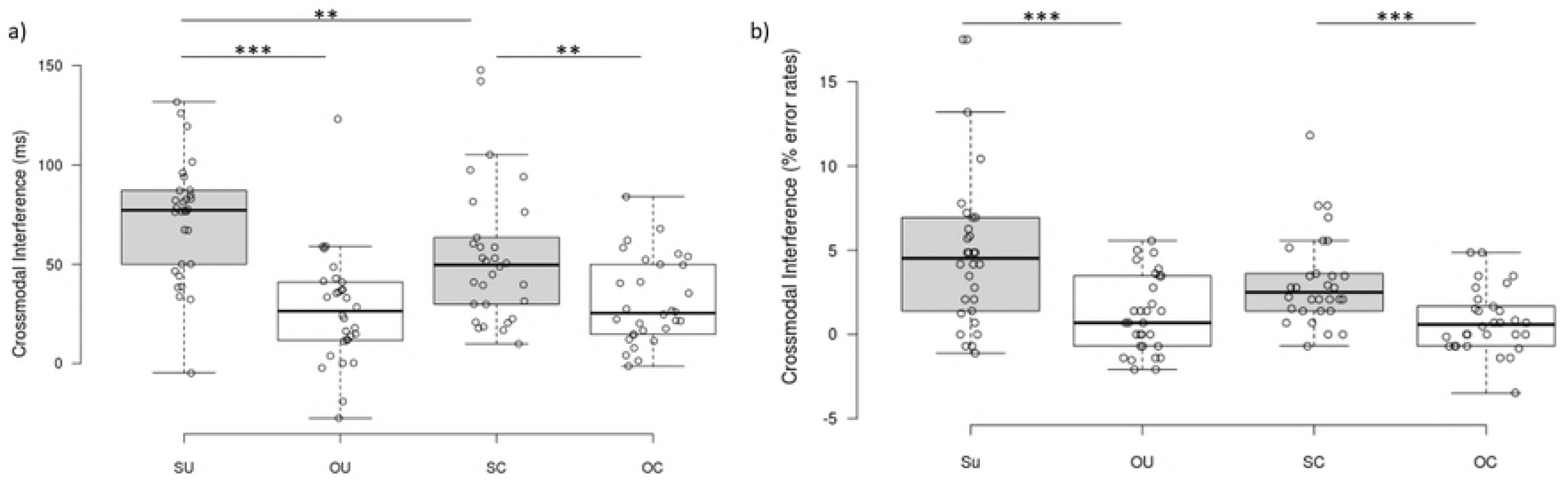
Crossmodal interference – three-way interaction. Crossmodal interference shown by Tool Arrangement (uncrossed [U], crossed [C]) and Visual Field (same [S], opposite [O]) for reaction times (A) and percentage error rates (B), on the Crossmodal Congruency Task (CCT), for all participants (n = 30). Crossmodal interference was calculated by subtracting congruent from incongruent reaction times and error rates. Medians are depicted by the centre lines. The box limits indicate the 25^th^ and 75^th^ percentile. The whiskers extend 1.5 times the interquartile range from the box limits. Circles depict individual data points. ** *p* < .01, *** *p* < .001.

A four-way interaction between the factors Set, Tool Arrangement, Visual Field and Congruence on RTs for the CCT was also observed (Fig 4), *F*(2.41, 70.09) = 3.28, *p* = .035, η^2^_p_ = .10. Separate threeway ANOVAs of Tool Arrangement x Visual Field x Congruence for each Set revealed significant three-way interaction for only the first two sets (set 1 [passive]: F(1, 29) = 8.89, *p* = .006, η^2^_p_ = .24; set 2: F(1, 29) = 11.09, *p* = .002, η^2^_p_ = .28). This interaction was not present in set 3, F(1, 29) = 0.39, *p* = .536, η^2^_p_ = .01, or set 4, *F*(1, 29) = 0.11, *p* = .748, η^2^_p_ < .01. To further investigate these patterns of results we calculated the crossmodal interference and compared this across each level of Tool Arrangement and Visual Field with each set (Fig 4).

**Fig 4.**
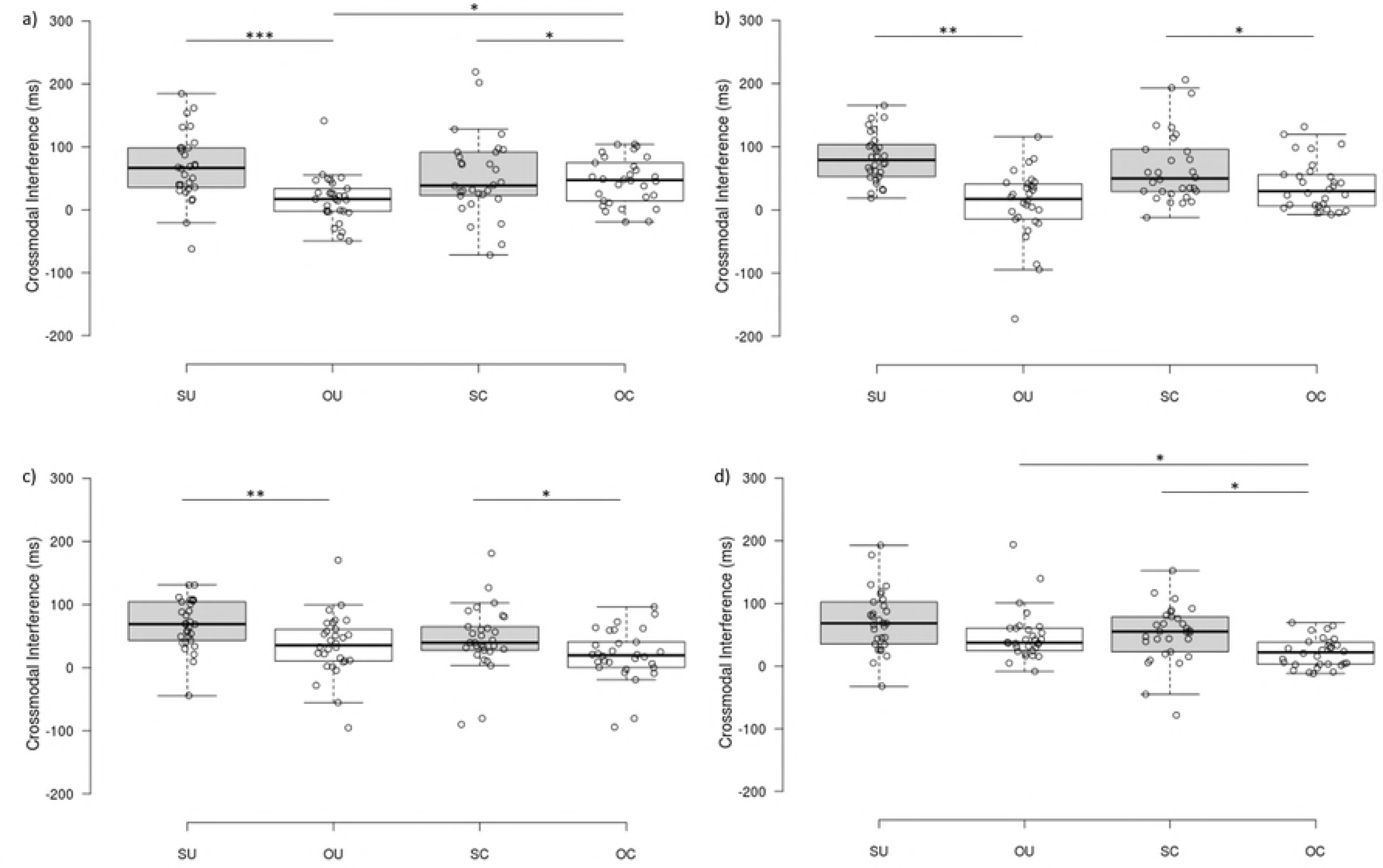
Crossmodal interference – four-way interaction. Crossmodal interference shown by Set (1 [passive], 2, 3, 4) Tool Arrangement (uncrossed, crossed) and Visual Field (same, opposite) for reaction times on the Crossmodal Congruency Task (CCT), for all participants (n = 30). Crossmodal interference was calculated by subtracting congruent from incongruent reaction times. Circles depict individual data points. Medians are depicted by the centre lines. The box limits indicate the 25ht and 75^th^ percentile. The whiskers extend 1.5 times the interquartile range from the box limits. Circles depict individual data points * *p* < .05, ** *p* < .01, *** *p* < .001.

Follow-up analysis (see S3 Table) of the crossmodal interference scores showed significantly greater interference for same side than opposite side distractors for uncrossed tools for set 1 (passive), *t*(29) = 4.84, *p*_adjusted_ < .001, *d* = 1.80, set 2, *t*(29) = 5.59, *p*_adjusted_ = .004, *d* = 2.08, and set 3, *t*(29) = 3.77, *p*_adjusted_ = .004, *d* = 1.40, but not for set 4, *t*(29) = 1.98, *p*_adjusted_ = .124, *d* = 0.74. Crossmodal interference was also greater for same side compared to opposite side distractors when tools were crossed for set 1 (passive), *t*(29) = 2.76, *p*_adjusted_ = .042, *d* = 1.03, set 2, *t*(29) = 2.74, *p*_adjusted_ = .015, *d* = 1.00, set 3, *t*(29) = 2.45, *p*_adjusted_ = .048, *d* = 1.00, and set 4, *t*(29) = 2.84, *p*_adjusted_ = .012, *d* = 1.05. For visual distractors appearing in the opposite visual field relative to vibrotactile targets, crossmodal interference was greater for crossed than uncrossed tools for set 1, *t*(29) = 2.78, *p*_adjusted_ = .042, *d* = 1.03. There were no significant differences for opposite visual field distractors between crossed and uncrossed tools for sets 2 and 3 *t*s(29) ≤ 2.30, *ps*_adjusted_ ≥ .058, *ds* ≤ 0.85. For set 4, however, crossmodal interference was greater for uncrossed than crossed tools, when visual distractors appeared in the opposite visual field relative to vibrotactile targets, *t*(29) = 3.09, *p*_adjusted_ = .045, *d* = 15. There were no significant differences for visual distractors appearing in the same visual field relative to vibrotactile targets, between crossed and uncrossed tools, *t*s(29) ≤ 1.71, *ps*_adjusted_ ≥ .138, *ds* = 0.64. Overall, these results show that the expected pattern of differences in interference between crossed and uncrossed conditions, reflecting that the tool tips were incorporated into peripersonal space, was evident in sets 1 (passive), 2, and 3. That is, the magnitude of crossmodal interference for distractors in the same compared to opposite visual field was smaller when tools were crossed compared to uncrossed. However, our results show that this pattern of crossmodal interference was reversed in set 4. Overall, the change in crossmodal interference across the four sets of the CCT task is not consistent with a gradual emergence of the effects of tool-use on peripersonal space over time.

### Tactile distance judgements

There was a significant main effect of Set, reflecting a decrease in TDJ from pre (*M* = 9.67, *SD* = 3.19) to post active tool-use (*M* = 10.09, *SD* = 3.36), when using a one-tailed test on an *a priori* basis, *F*(1, 29) = 3.20, *p* = .041, η^2^_p_ = .10. There were no other main effects or interactions for the TDJ, including none involving Sensory Condition, *Fs*(1, 29) ≤ 2.69, *ps* ≥ .111, η^2^_p_s ≤ .09.

### Exploratory analyses

The above results show that the pattern of interference during set 1 of the CCT is consistent with updating of peripersonal space (Fig 4a). This was unexpected, given that set 1 required only passive interaction with the tools. We considered that this could be due to the repeated-measures design of the study. That is, experience with the tool in session 1 might have primed participants to rapidly embody the tools upon grasping the handles of the tools at the beginning of sessions 2 and 3, extending peripersonal space even while passively interacting with the tools. Because the order of the study was randomised and counterbalanced, we could investigate this possibility by conducting a between groups analysis of the CCT data from only the first session, when there was no prior experience with the tools. That is, in this exploratory analysis Sensory Condition was treated as a between-subjects factor with ten participants in each of the pain, active placebo, and neutral groups. We conducted two five-way ANOVAs on the RTs and error rates from the first experimental session with Set, Side of Body, Tool Arrangement, Visual Field and Congruence as within-subjects factors; and Sensory Condition as a between-subjects factor. There was no main effect of Sensory Condition on RTs from the CCT, *F*(2, 27) = 0.97, *p* = .390, η^2^_p_ = .07. There was no clear effect of Sensory Condition or any interactions of interest for error rates from the CCT during session 1 (see S1 Text). Furthermore, when the session order was included as a variable in the main analysis there was no change to the key interaction terms. Therefore, it seems unlikely that the apparent extension of peripersonal space during the first set of the CCT can be attributed to familiarity with the tools due to the repeated-measures design of the study.

To explore the evidence for the null hypothesis, we reanalysed the main interaction terms from the CCT and TDJs that involved Sensory Condition with a Bayesian repeated-measures ANOVA using JASP software [67]. To calculate the adjusted BF_10_, we divided the posterior probability of the models, or P(M|data), from the model that included the interaction term of interest, by the model containing all other elements of the first model except from the interaction term of interest [68]. We found moderate evidence [69] of no effect of an interaction between Sensory Condition, Tool Arrangement, Visual Field, and Congruence on RTs from the CCT (adjusted BF_10_ = 0.12). We also found moderate evidence of no effect of an interaction between Sensory Condition and Set on TDJs (adjusted BF_10_ = 0.24). These findings support our interpretation that pain induction did not interfere with the updating of peripersonal space and body representations.

## Discussion

Our study aimed to investigate the effect of induced pain on updating of peripersonal space and body representations during and following tool-use. We hypothesised that participants would be less able to update peripersonal space and body representations during pain induction to the arm, compared to the two control conditions. We used a crossmodal congruency task (CCT) and tactile distance judgements (TDJs) to measure updating of peripersonal space and body representations, respectively. The global patterns of the CCT and TDJ were consistent with previously reported effects of tool-use (e.g. [48, 50]). In contrast to our predictions, we found that pain did not interfere with updating of peripersonal space and body representations following active tool-use, when compared to two control conditions (i.e. active placebo, and neutral). That is, we found evidence that the performance on the CCT and TDJ did not differ between sensory conditions. There was also no significant difference in the CCT or TDJ when we explored pain ratings as a covariate. Therefore, experimentally induced pain does not appear to influence the updating of peripersonal space and body representations during and following tool-use.

It is unlikely that the lack of an effect of pain can be attributed to failure of our protocols to induce updating in peripersonal space and body representation. For reaction times from the CCT, we found that reaction times to vibrotactile targets were slower when accompanied by visual distractors in the same visual field compared to the opposite, but this effect was weaker when tools were crossed such that the opposite side visual distractors appeared on the same tools as the vibrotactile targets. These findings are comparable to the results reported by Maravita and colleagues [48]. We also found that estimates of the felt distance between two points (TDJs) parallel to the axis of the tool decreased in both arms after active tool-use. This is thought to reflect that the body representation has updated to incorporate the tools, and is consistent with previous findings (e.g.[50]). Our study thus replicated evidence of updating peripersonal space and body representations during and after tool-use, but induced pain did not modulate these effects.

It is also unlikely that the absence of a significant effect of pain on peripersonal space and body representation in this study is due to failure of our sensory manipulations or compensatory changes in movements during pain induction. Participants reported experiencing pain throughout the study in the pain condition and not for the two other conditions, indicating that our pain induction was successful. We confirmed that movement patterns were similar for all three conditions by having a condition-blind observer rate videos of participants’ movements. We also found that mechanical pain threshold (MPT) on the finger increased after the pain induction to the arm, and this change in MPT remained until the end of the study. This demonstrates that our manipulation altered sensory processing relevant to the hand. However, mechanical detection thresholds remained unchanged, indicating that the ability to detect a tactile stimulation was the same across sensory conditions. Therefore, our manipulation succeeded in inducing pain, without impairing movement or tactile sensation, and so it is unlikely that our results can be attributed to methodological limitations.

Bodily and spatial representations can be influenced by pain. Previous work has demonstrated that spatial perception can be modified by experimentally induced pain. For instance, the subjective body midline deviated towards a painful thermal stimulation with a large magnitude of effect [46], and painful cooling can increase the felt size of the thumb [47]. Our results, however, suggest that pain does not alter the flexibility of spatial and body representations to *update* as a result of interaction with objects in our environment (e.g. during tool-use). This has ramifications for how distortions in body representation and peripersonal space are maintained in people with chronic pain. Specifically, it could suggest that pain might not be the driving factor preventing normal body representation and peripersonal space from being restored.

An alternative perspective on our results from the CCT, showing no effect of pain, might be offered by the distinction between goal-directed and defensive dimensions of peripersonal space, as proposed by De Vignemont and Iannetti [10]. They conceptualise goal-directed peripersonal space as the space upon which we can act, and defensive peripersonal space as the space in which we might have to react to incoming, and potentially harmful, objects. Research into tool-use largely covers goal-directed movements and tasks, and so Vignemont and Iannetti [10] speculated that defensive space would not be modulated by tool-use. It could be that the painful stimulation used in our study altered properties of defensive peripersonal space, whereas our task measured changes in goal-directed peripersonal space representations. Although they serve separate functions, there is evidence to suggest that goal-directed and defensive peripersonal space representations can interact. Rossetti and colleagues [70] showed that incoming painful stimuli, in this case a 4 cm long medical needle, presented both at 20 and 40 cm away from the body triggered an alerting response (as measured by skin conductance response) in healthy participants, but only after active use of a 40 cm tool. This shows that tool-use can modulate a response to an incoming painful stimulus. Our study, however, shows that acute pain does not alter the updating of goal-directed peripersonal space. More research is needed to characterise how goal-directed and defensive peripersonal space representations interact, and how different qualities of pain might influence such interactions. For instance, it could be that acute pain alters defensive peripersonal space in healthy individuals, as is the case in in people with trigeminal neuralgia [71], or that modifications of defensive peripersonal space are limited to approaching painful stimuli (i.e. when there is the potential threat of pain).

Although our findings were qualitatively similar to Maravita and colleagues [48] in that we found overall less interference from opposite-side distractors when the tools were crossed compared to uncrossed, these differences were less pronounced in our study. This was despite the fact that we included additional interactive tool use (beanbag sorting) tasks between the three active sets of the CCT task. We also did not replicate the expected effect of active tool-use on performance on the CCT over time. That is, we did not find that interference effects thought to reflect expansion of peripersonal space increased over time. Instead, we observed a decrease in this pattern as participants spent more time interacting with the tool. Furthermore, we found that participants showed interference effects consistent with expansion of peripersonal space during passive interaction with the tools. It is unclear why our results differ from those reported by Maravita and colleagues [48]. Although our CCT task replicated that of their study in most respects, a key difference is that we asked participants to indicate the location of the vibrotactile targets using a four-alternative forced choice response (the factorial combination of up-down *and* left-right). Maravita and colleges [48] used a two-alternate forced choice response in which participants indicated only the up-down location of the vibrotactile stimuli regardless of the side of space upon which they were presented. We used the four-alternative forced-choice response because we sought to disentangle limb-specific effects of unilateral pain induction. That is, if we had found that pain interfered with updating of peripersonal space and body representations, we aimed to explore whether this interference was restricted to the side of the painful arm, or if pain disrupted these processes more generally. It is possible that our four-alternative forced-choice response added an additional level of spatial incongruence that prevented the emergence of a stronger effect of tool use in this task, as the crossmodal congruency effect is driven by the reaction time cost that arises from presenting visual distractors at spatially incongruent locations to tactile targets [72]. For example, in our study spatial incongruence could be created when the tools were crossed and the distractor originated on the same tool and in the same vertical location as the vibrotactile target (i.e. the distractor and target are in opposite visual fields). In the study of Maravita and colleagues [48], however, no such spatial incongruence would have been present in such a trial with regards to the response required (up or down), thus making object-based effects easier to interpret. This might explain why we found an overall pattern that was comparable to Maravita and colleagues, indicative of peripersonal space updating, although the effect was less pronounced. Future studies should limit themselves to one level of spatial incongruence (e.g. up/down responses only).

To our knowledge, this was the first study testing changes in TDJs in both arms (rather than just one) following tool use. Previous studies using this method have tested only one arm, however it is conceivable that there could be differences in how body representation is updated in the two arms, for example due to differences in activity levels between arms (e.g. [73]). Our results showed no difference between the change in TDJs for the two arms.

To conclude, we sought to investigate the effect of induced pain on the updating of peripersonal space and body representations during and following tool-use. Our study replicated findings showing that active tool-use updated peripersonal space and body representations. We also successfully induced pain, without impairing movement or tactile sensitivity. However, we found evidence that induced pain did not interfere with *updating* peripersonal space and body representations. When considered with previous results, these results suggest that induced pain can cause a direct change in bodily and spatial perception, but the mechanisms involved in *updating* such representations do not appear to be disrupted. This suggests that any disruption to these processes in pathological pain conditions cannot be sufficiently explained by acute pain.

## Supporting information

**S1 Fig. Example tool arrangement**. The different possible combinations for the Crossmodal Congruency task (CCT) for congruent visuotactile stimulation. The brighter red dots illustrate the location of the visual stimulation, and the yellow stars the location of the tactile stimulation for each example trial. The tools could be in the straight (A & B), or crossed (C & D) position, with light appearing in the same (A & C) or opposite (B & D) visual field as the visuotactile stimulation.

**S2 Fig. Pain ratings over time**. Change in pain ratings over time in the pain (red line) and active placebo (blue line) conditions. The neutral condition was omitted as mean ratings were ≈ 0. The ramp up period (t1-t15) lasted 15 min for the active placebo and neutral conditions. For the pain condition, this period lasted until pain ratings reached 5/10, or plateaued at 3/10 or higher for 3 consecutive ratings, which took on average 16.7 minutes (*SD* = 1.88). Pain ratings were recorded every minute during the ramp up period. When the ramp up period was shorter than 15 minutes, the missing ratings were adjusted to 5/10 for the purpose of this figure. *Pain ratings for ramp up periods exceeding 15 minutes are not included in this figure*. During the main experiment (t16-t27) pain ratings were recorded between the tactile distance judgements and different set of the tool-use tasks, and so the time between pain ratings was variable.

**S3 Fig. Change in Mechanical Pain Threshold by Sensory Condition**. Mechanical Pain Threshold (MPT) in mN on the middle finger (D3) before and after pain induction, active placebo, or no sensory manipulation (neutral), split by Side of Body for each conditions: neutral (A), active placebo (B), and pain (C). Circles depict individual data points. The dominant arm was always stimulated. (N = 30), * *p* < .05, ** *p* < .01.

**S4 Fig. Crossmodal interference – four-way interaction for error rates**. Crossmodal interference shown by Set (1 [passive], 2, 3, 4), Tool Arrangement (uncrossed, crossed), and Visual Field (same, opposite) for error rates expressed in percentages on the Crossmodal Congruency Task (CCT), for all participants (n = 30). Crossmodal interference was calculated by subtracting congruent from incongruent error rates. Medians are depicted by the centre lines. The box limits indicate the 25ht and 75^th^ percentile. The whiskers extend 1.5 times the interquartile range from the box limits. Circles depict individual data points. (N = 30). ** *p* < .010, *** *p* < .001.

**S1 Table. Performance on sensory tests by Sensory Condition**. Performance on sensory tests (Mechanical pain Threshold [MPT], Mechanical Detection Threshold [MDT], Two Point Discrimination [TPD]) expressed as change from pre to post Sensory Manipulation. Means and standard deviations for Sensory tests are split by Sensory Condition (Pain, Active Placebo, Natural), and Side of Body (dominant [stimulated], non-dominant).

**S2 Table. CCT main effects and interactions for reaction times**. All significant main effects interactions from six-way ANOVA of Sensory Condition, Side of Body, Set, Tool Arrangement, Visual Field, and Congruence from the Crossmodal Congruency Task for reaction times.

**S3 Table. CCT reaction times**. Descriptive statistics presented for the reaction times in ms from the crossmodal congruency task, split by Tool Arrangement (uncrossed, crossed), Side of Body (same, opposite), and Congruence (congruent, incongruent). The difference between congruent and incongruent scores are reported within each level of Tool Arrangement, and Side of Body.

**S4 Table. CCT main effects and interactions for error rates**. All significant main effects interactions from six-way ANOVA of Sensory Condition, Side of Body, Set, Tool Arrangement, Visual Field, and Congruence from the Crossmodal congruency task for error rates.

**S1 Text. Crossmodal congruency task (CCT) results – error rates**.

